# Neural signatures of model-free learning when avoiding harm to self and other

**DOI:** 10.1101/718106

**Authors:** Patricia L. Lockwood, Miriam Klein-Flügge, Ayat Abdurahman, Molly J. Crockett

**Affiliations:** Department of Experimental Psychology, University of Oxford, Oxford OX1 3PH, United Kingdom; Wellcome Centre for Integrative Neuroimaging, Department of Experimental Psychology, University of Oxford; Department of Psychology, Yale University, New Haven, CT, 06511, USA

**Author notes:** Correspondence should be addressed to: Patricia L. Lockwood, Experimental Psychology, Tinsley Building, University of Oxford, OX1 3SR, United Kingdom., Miriam C Klein-Flügge, Experimental Psychology, Tinsley Building, University of Oxford, OX1 3SR, United Kingdom., Molly J. Crockett, Department of Psychology, 2 Hillhouse Avenue, New Haven, CT 06511, USA. Indicates joint first authorship (Patricia L. Lockwood, Miriam Klein-Flügge).

## Abstract

Moral behaviour requires learning how our actions help or harm others. Theoretical accounts of learning propose a key division between ‘model-free’ algorithms that efficiently cache outcome values in actions and ‘model-based’ algorithms that prospectively map actions to outcomes, a distinction that may be critical for moral learning. Here, we tested the engagement of these learning mechanisms and their neural basis as participants learned to avoid painful electric shocks for themselves and a stranger. We found that model-free learning was prioritized when avoiding harm to others compared to oneself. Model-free prediction errors for others relative to self were tracked in the thalamus/caudate at the time of the outcome. At the time of choice, a signature of model-free moral learning was associated with responses in subgenual anterior cingulate cortex (sgACC), and resisting this model-free influence was predicted by stronger connectivity between sgACC and dorsolateral prefrontal cortex. Finally, multiple behavioural and neural correlates of model-free moral learning varied with individual differences in moral judgment. Our findings suggest moral learning favours efficiency over flexibility and is underpinned by specific neural mechanisms.

## Introduction

A central component of human morality is a prohibition against harming others^1,2^. People readily avoid actions that might harm another person^3–7^, and this basic harm aversion is so strong that many people even find it distressing to perform pretend harmful actions, such as shooting someone with a fake gun^8^. Harm aversion is disrupted in clinical disorders such as psychopathy that have a strong developmental component^9^, and although harm aversion is robust in healthy adults, anyone who has watched young children fighting over a coveted toy knows that such an aversion is not present from birth. Indeed, a large literature documents the emergence of moral conduct over the course of development^10,11^. Cross-cultural differences in morality suggest moral behavior is fine-tuned to local environmental demands^12^, and lab experiments demonstrate how individuals can quickly adapt moral behavior to changing norms^13,14^. All this evidence highlights a critical role for *learning* in the development of harm aversion and moral behaviour more broadly^6^.

Recent work in computational neuroscience has advanced our knowledge of how organisms learn the value of actions and outcomes via reward and punishment^15,16^. An important theoretical distinction has been made between ‘model-based’ and ‘model-free’ learning^17,18^. The computationally expensive model-based system builds an internal model of the environment and selects actions by prospectively searching the internal model for the best course of action^19,20^. The computationally efficient model-free system assigns values to actions through trial-and-error and selects actions retrospectively based on these ‘cached’ values; because it does not store a model of the environment, it can sometimes make sub-optimal recommendations (e.g., in settings where typically valuable actions lead to bad outcomes, or vice versa). These two systems are somewhat neurally dissociable, with model-based learning preferentially engaging lateral prefrontal cortex (LPFC), posterior parietal cortex and caudate^20–22^ and model-free learning preferentially engaging putamen^23,24^, although both systems update their representations via prediction errors encoded in overlapping regions of ventral striatum^20^. Model-based and model-free systems often make similar recommendations about which actions are more valuable, but when they conflict, an arbitration process allocates control between them^22,25,26^. However, despite extensive theorizing that the model-based/model-free distinction may help to characterize puzzling features of moral learning and decision-making^3,27–29^, it remains unknown whether the moral consequences of actions affect the balance between model-based and model-free control, and whether common or distinct neural processes are engaged when learning to avoid harmful outcomes to self and others.

Past work on the neural basis of moral decision-making provides support for competing hypotheses. On the one hand, the sophistication of human morality seems to demand the kinds of complex representations afforded by model-based learning, suggesting learning to avoid harming others may preferentially engage the model-based system. Supporting this view, people are easily able to learn to avoid harmful actions without directly experiencing their outcomes through trial and error, as may be required for model-based learning^3,27,30^. Moreover, moral decision-making in healthy adults consistently engages brain regions most strongly associated with the model-based system, including lateral prefrontal cortex (LPFC), caudate, and temporoparietal junction (TPJ)^23,25,31^. Deciding to follow moral norms like fairness and honesty, and enforcing those norms on others via costly punishment, engages LPFC^32–37^, and disrupting LPFC function reduces moral norm compliance and enforcement^38,39^. During decisions to avoid harming others, LPFC encodes the blameworthiness of harmful choices and modulates action values in caudate and thalamus^4^, two subcortical areas shown to play a critical role in associative learning and pain processing as well as moral decision-making^40–45^. TPJ has been implicated in sophisticated representations of others’ mental states and integrating these into social decisions^46–48^.

On the other hand, one principal function of model-free learning is to cache value in actions that are reliably adaptive, sacrificing flexibility for efficiency. Given that harming others is typically prohibited, actions that harm others may represent a special class of actions that are prioritized for model-free learning, similar to how certain classes of stimuli, like snakes and spiders, are “prepared” for aversive classical conditioning^49^. In other words, since avoiding harm to others is hugely important for social life, learning processes that fast-track harm-avoidant action selection to a habitual, automatic process may be socially adaptive. Supporting this view, recent work suggests that morality constrains mental representations of what actions are considered possible; harmful actions are removed from choice sets as a default^50^, and choices that harm others are slower than helpful choices, suggesting an automatic tendency to avoid harm^5,51,52^. Furthermore, recent studies of model-free learning to gain rewards for oneself and others have highlighted a distinct encoding of prediction errors concerning others’ outcomes in the subgenual anterior cingulate cortex (sgACC)^53,54^, a region that has been implicated in social and moral decision-making more broadly^54–58^. Model-free processes that distinguish learning about how one’s actions affect others could provide a neural mechanism for prioritizing model-free learning in moral contexts.

To test these competing hypotheses, we used computational modelling and fMRI to probe the relative balance between model-based versus model-free processes, and their neural bases, when people learn to avoid moderately painful electric shocks for themselves and a stranger. After undergoing a pain thresholding procedure (see Methods), participants (N=36) completed a hybrid version of two paradigms previously proposed to reliably dissociate model-free versus model-based learning (Figure 1)^20,22,31^. We optimized the task in a way that allowed us to address the specific hypotheses examined in the present study (see Supplementary Note on Experimental Design for details).

**Figure 1.**
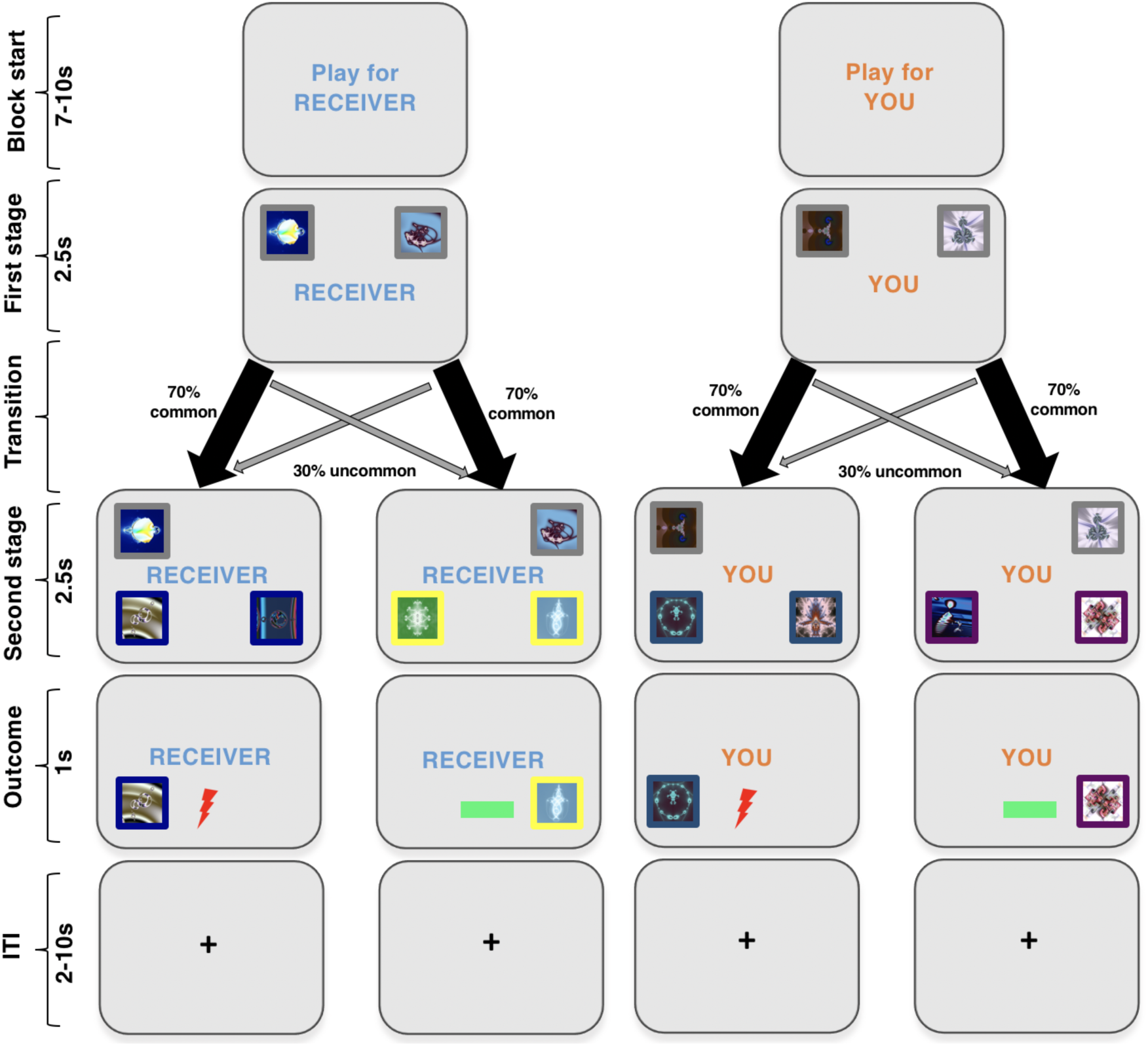
Model-free and model-based aversive learning task. Participants completed a two-stage decision-making task to assess the tendency to engage in model-free and model-based learning. The task was a hybrid of two tasks previously shown to assess model-free and model-based learning processes^20,26^ to probe learning to avoid aversive (shock) outcomes for either oneself or another person (the ‘receiver’, referred to as ‘other’ hereafter). At the beginning of each block an instruction cue signalled the recipient of the outcome (self or other). At the first stage two images were displayed that probabilistically led to one of two states depicted by different colours surrounding the boxes. In this example, to ‘blue zone’ or ‘yellow zone’ for the other participant, and ‘turquoise zone’ or ‘purple zone’ for self. Participants then made a second choice between two pictures in the coloured zone which was followed by an outcome of shock or no shock. The probability with which the boxes at the second stage delivered a shock or no-shock outcome drifted throughout the experiment (bounded between 0% and 100% with a drift rate of 0.2) and participants were instructed to keep learning throughout. 10% of the total electric shocks accumulated in the ‘self’ condition were delivered to the participant themselves at the end of the experiment whilst 10% of the electric shocks accumulated in the ‘other’ condition were delivered to the partner participant.

## Results

### Model-free learning is prioritized when avoiding harm to others vs. self

Participants completed a two-step decision-making task to index model-free and model-based learning strategies (**Figure 1**). Prior to scanning, participants were trained on the transition structure of the task using stimuli different from the main experiment which allowed them to learn the probabilistic transition structure. The two-step task distinguishes model-free and model-based learning by measuring people’s choices to stay or switch based on the outcome of the previous trial and the transition structure. Theoretically, a purely model-free learner would ignore the transition structure and repeat first-stage choices if they prevented pain on the previous trial but switch choices if the previous choice caused pain. Thus, model-free learning is reflected in a main effect of outcome (pain versus no pain) on subsequent first-stage choice behaviour. In contrast, a model-based learner would take the transition structure into account. While behaviour on common transitions would be similar for a model-free agent, after rare transitions, a model-based agent would repeat a first-stage choice if the outcome was pain, but switch if the outcome was no pain. Thus, model-based learning is reflected in an interaction between outcome (pain versus no pain) and transition (common versus rare). It is now well established that people display a combination of model-free and model-based learning strategies when learning about rewarding outcomes^20,31,59^ and initial evidence indicates that the same is true for learning about aversive outcomes for oneself^60,61^. We therefore first examined whether participants displayed a combination of model-free and model-based processes during aversive learning for oneself and others, as observed in these previous studies.

We performed a logistic regression analysis predicting first-stage stay versus switch choices as a function of the outcome (pain or no pain), transition on the previous trial (common or rare) and recipient (Self vs. Other). We included all main effects and interactions in the model. We found a significant main effect of outcome (*t*(35)=4.618, *p*<.001, confidence interval (CI) for beta estimate: 0.17, 0.42, Cohen’s *d* = 0.77) indicating a contribution of model-free learning, but also a significant transition by outcome interaction (*t*(35)=-3.173, *p*=.003, CI for beta estimate: − 0.23, − 0.05, *d* = −0.53), indicating the presence of model-based learning (**Figure 2a**,**b**). This demonstrates that similar to reward learning, aversive learning is underpinned by a mixture of model-based and model-free processes.

**Figure 2.**
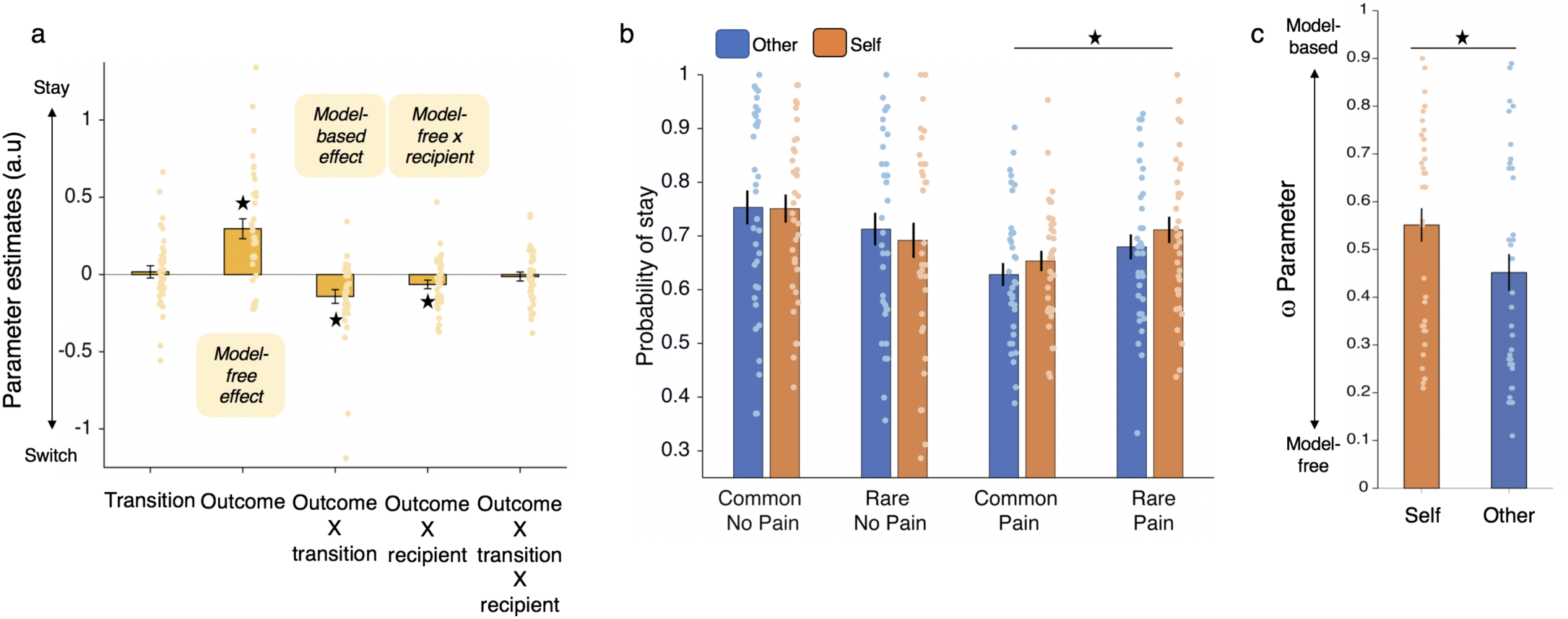
Model-free and model-based choices when avoiding harming oneself and others. (a) Logistic regression coefficients predicting first-level stay/switch choices (mean ± SEM). Participants exhibited a main effect of staying after no pain which indicated model free behaviour (*t*(35)=4.618, *p*<.001, confidence interval (CI) for beta estimate: 0.17, 0.42, Cohen’s *d* = 0.77) and a outcome by transition interaction which indicated model-based behaviour (*t*(35)=-3.173, *p*=.003, CI for beta estimate: −0.23, −0.05, *d* = −0.53). Intriguingly there was also a outcome x recipient interaction ((*t*(35)=- 2.31, *p*=.027, CI of beta estimates −0.12, −0.008, *d* = −0.39) showing that participants were more model-free, and thus more likely to switch after pain and stay after no pain, independent of transition type, when making choices for another person. (b). The probability of repeating a choice at the first-level (‘stay’) is plotted as a function of the transition and outcome on the previous trial. This shows that the outcome x recipient interaction in (a) is mostly driven by fewer stay trials after pain, regardless of transition (two rightmost blue vs orange bars). Thus, the more pronounced model free behaviour for others is mostly driven by a lower probability of staying after pain outcomes rather than a higher probability of staying after no pain outcomes). (c) ω estimates from the best fitting model showed that the ω parameter was significantly lower for other (0.45) than self (0.55) consistent with the regression analyses that showed people were more model free when avoiding harm to others compared to self (*p*<.02). Asterisks indicate significant difference at *p*<.05.

**Figure 3.**
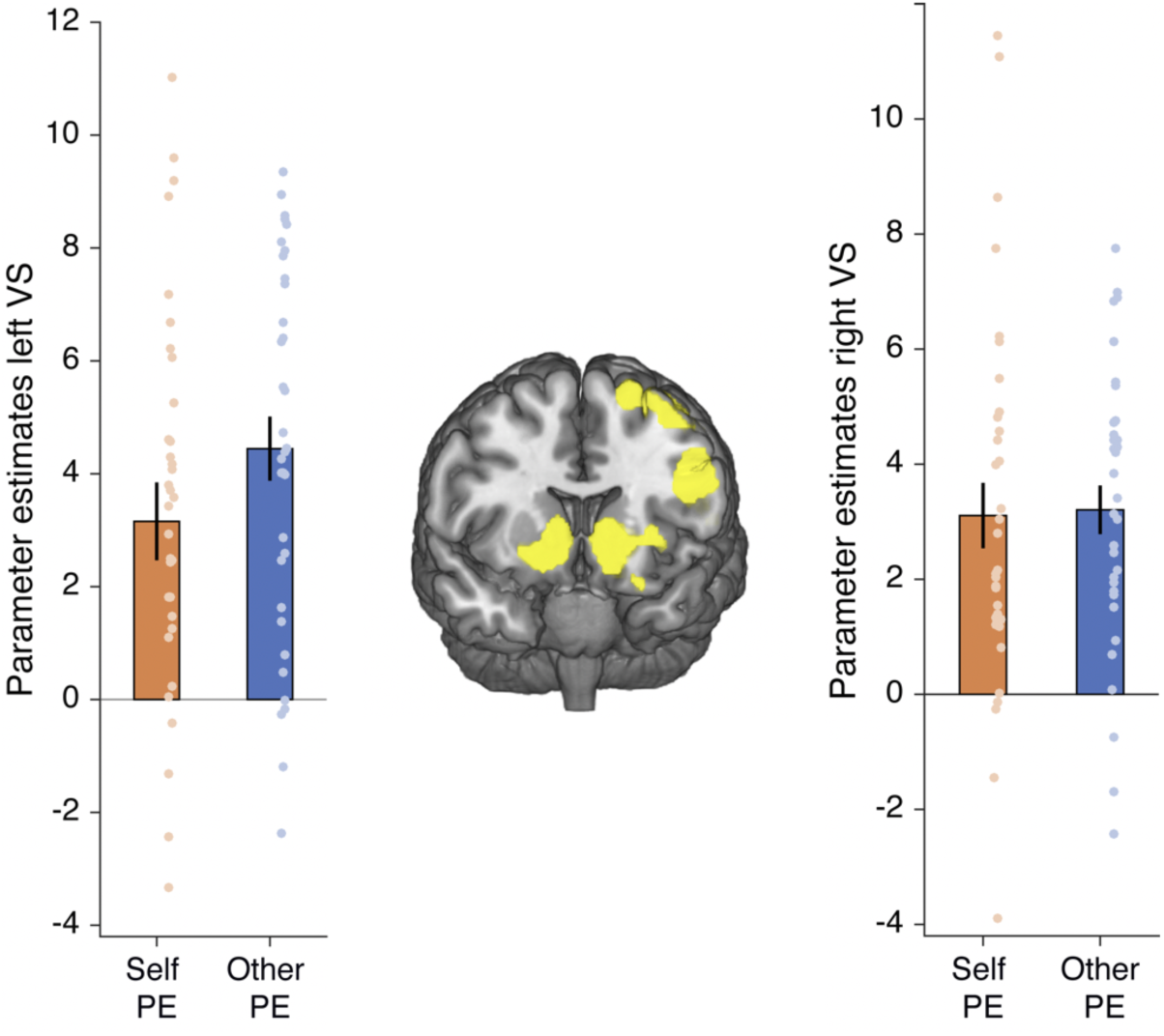
Ventral striatum encodes prediction errors of pain avoidance for self and other. Ventral striatum (right x = 10, y = 12, z = −4, k=236, z= 5.84, left x = −16, y = 6, z = −10, k=458, z= 5.77, *p*<.05 FWE-whole brain corrected after initial thresholding at *p*<.001) tracked model-free prediction errors for both self and other bilaterally, with no significant differences between conditions.

Intriguingly, we also found a significant interaction between outcome (the model-free contribution to learning) and recipient, showing that people were more model-free for others relative to self (*t*(35)=-2.31, *p*=.027, CI of beta estimates −0.12, −0.008, *d* = − 0.39). By contrast, there was no interaction between recipient and transition x outcome, the model-based component of learning (*t*(35)=-.459, *p*>.65, CI of beta estimates: −0.071, 0.045, *d* = −0.08).

We validated these behavioural results by repeating our analysis using the lme4 package in R which ensured we had good estimates of random effects and accounted for variability in behaviour using Bound Optimization by Quadratic Approximation (see Methods). Our model predicted the tendency to switch vs. stay including all main effects and interactions and a random effect of subject, glmer(Switch_Stay∼NoPain_Pain*Transition*Agent+(1|Subject). All results remained the same (main effect of outcome (model-free) Z=5.949, *p*<.001; Outcome x transition interaction (model-based) Z=5.484, *p*<.001; outcome x recipient Z=-2.154, *p*=0.031; [outcome x transition (model-based)] x recipient Z=0.696, *p*=0.4864). Again, these findings support the idea that people were more model-free when avoiding harming others.

To further examine which outcomes most influenced the outcome x recipient interaction, we performed separate post-hoc tests on the percentage of stay/switch choices for self and other following (a) only pain outcomes and (b) only no pain outcomes. This showed that the recipient difference was driven by increased switching after pain outcomes for other versus self (*p*=.028), with no difference between recipients in the proportion of stay/switch choices after no pain outcomes (*p*=.552). This is important because the specificity of the effect rules out that people were simply more indifferent or inattentive to the outcomes of others compared to self.

### Computational modelling of aversive learning for self and other

Next, we fitted several trial-by-trial computational models to our data to examine further which model best captured the described behaviours during aversive learning for self and other. Deriving such trial-by-trial estimates that capture individual choice preferences was a prerequisite for modelling the fMRI data and allowed us to support our logistic regression analyses. We started with the full seven-parameter model proposed by Daw and colleagues^20^ and compared this model to similar models with fewer parameters (four of five) following modifications similar to those suggested in previous studies (e.g.^62,63^); for details, see Methods and **Supplementary Table 2**). We also included variants of the same models that involved separate learning rates for pain and no pain outcomes, given evidence suggesting differential learning as a function of outcome valence (e.g.^64,65^). All of these models were initially fitted separately on self and other blocks.

We found that a five-parameter model best explained behaviour compared to all alternative models tested. This model included separate learning rates for no pain and pain outcomes (αPain, αNoPain), a single temperature parameter capturing choice randomness (β), a perseverance parameter capturing a tendency to stick with the previously made choice (ρ) and a model-free/model-based weighting parameter (ω). Importantly, this five-parameter model best explained behaviour in both the self and other blocks (**Supplementary Table 2**).

We next compared the different estimated parameters for the self and other blocks. This analysis showed a significant difference between the conditions in both the perseverance parameter ρ (*t*(35)=2.41, *p*=.02,) and the model-free/based weighting parameter ω (*t*(35)=3.10, *p*=.0039; **Supplementary Table 3**). To compare these parameters more robustly, we then used maximum a-posteriori estimation performed on the pooled data of self and other blocks to compare three models, one with separate perseverance ρ and ω parameters for self and other (and thus a total of seven parameters), one with separate ω parameters for self and other (and therefore a total of six parameters), and the original 5 parameter model (which assumes the same ρ and ω across self and other blocks). We used these models to examine whether differences in the ρ and ω parameters between self and other when fitted separately reflected true differences in the weighting of these parameters in a model comparison. This analysis showed that the model with separate ω’s for self and other, but not separate ρ’s, best explained the data. Importantly, these ω parameters were also significantly different from one another (selfω = 0.55, otherω = 0.45, *t*(35)=2.41, *p*=.02, d=0.40, **Figure 2c and Supplementary Table 4 and 5**). Thus, consistent with the regression based behavioural analyses that did not rely on a computational model (**Figure 2a**), participants were more model-free than model-based when learning to avoid harming others, compared to self (**Figure 2c**).

### Subcortical areas distinguish model-free prediction errors for self and other

Previous neuroimaging studies of model-based and model-free reward learning have reported model-free prediction error signals in ventral striatum^20^. We therefore first sought to replicate this effect in our aversive learning paradigm. To facilitate comparison with previous studies of reward learning, no-pain outcomes were coded as 1 and pain outcomes coded as −1. Therefore, a positive prediction error represents unexpected pain relief/avoidance, and a negative prediction error represents unexpected pain.

We built a general linear model (GLM1) that contained onsets for the first stage choice, second stage choice and outcome separately for self and other trials. These three time periods were each associated with parametric modulators from our winning model. These included the value difference between the two options at the first stage choice; the state prediction error based on the transition at the second stage choice; and the model-free prediction error at the time of the outcome. We focused our analysis on model-free prediction errors at the time of the outcome for two reasons. Firstly, our behavioural effects showed that self/other differences in learning emerged for model-free but not model-based learning. Secondly, model-free and model-based prediction errors are highly correlated and careful examination of their separate influences has shown that they are both encoded in ventral striatum^20^.

We began our analyses by examining whether previously reported neural correlates of value difference and state prediction errors were also observed in our novel paradigm. Several areas tracked inverse value difference and thus showed larger responses for choices that had a smaller value difference between the two first-stage options, including the most dorsal parts of anterior cingulate cortex near pre-SMA, bilateral inferior parietal cortex and middle frontal gyrus (**Supplementary table 1**), regions previously associated with the tracking of inverse subjective value difference^66–68^. These signals did not differ between self and other (see **Supplementary Table 1**). Also consistent with previous findings^21^, we found evidence of a main effect for state prediction errors at the second stage in dorsal ACC (x=-6, y=10, z=52, Z=4.85, K=906, *p*<.001 FWE-corrected) that again showed overlap between self and other (see **Supplementary table 1**).

Next, we tested whether model-free prediction errors were present in ventral striatum, as reported in a previous study examining model-based and model-free reward learning for self^20^ and several studies of reward based reinforcement learning^69^. We found a large bilateral cluster signalling prediction errors of harm avoidance (positive for no pain, negative for pain) in ventral striatum (right x = 10, y = 12, z = −4, k=236, Z= 5.84, left x = −16, y = 6, z = −10, k=458, z= 5.77, *p*<.05 FWE-whole brain corrected after initial thresholding at *p*<.001). Again, this signal did not significantly differ for self and other conditions (Right *t*(34) = −.14, *p*=.89, Left *t*(34) = −1.47, *p* = .152)).

Given that our behavioral results indicated model-free learning was prioritized when avoiding harm to others (relative to self), we next sought to identify areas that distinguished model-free prediction errors for others (relative to self). This analysis revealed a cluster in the thalamus extending into the caudate (x =16, y = −18, z =0, k=84, *p*=.033, Z=3.50 FWE-small volume corrected (SVC) after initial thresholding at *p*<.001, see Methods **Figure 4a**,**b**). The cluster positively tracked prediction errors of pain avoidance when learning for other (*t*(33)= 2.30, *p*=.028) and negatively tracked prediction errors of pain avoidance when learning for self (*t*(33)= −2.89, *p*=.007). Although this cluster extended into the caudate, the caudate ROI itself was not significant (x =18, y = 6, z =0, Z=3.49, k=2, *p*=.064, FWE-SVC after initial thresholding at *p*<.001).

**Figure 4.**
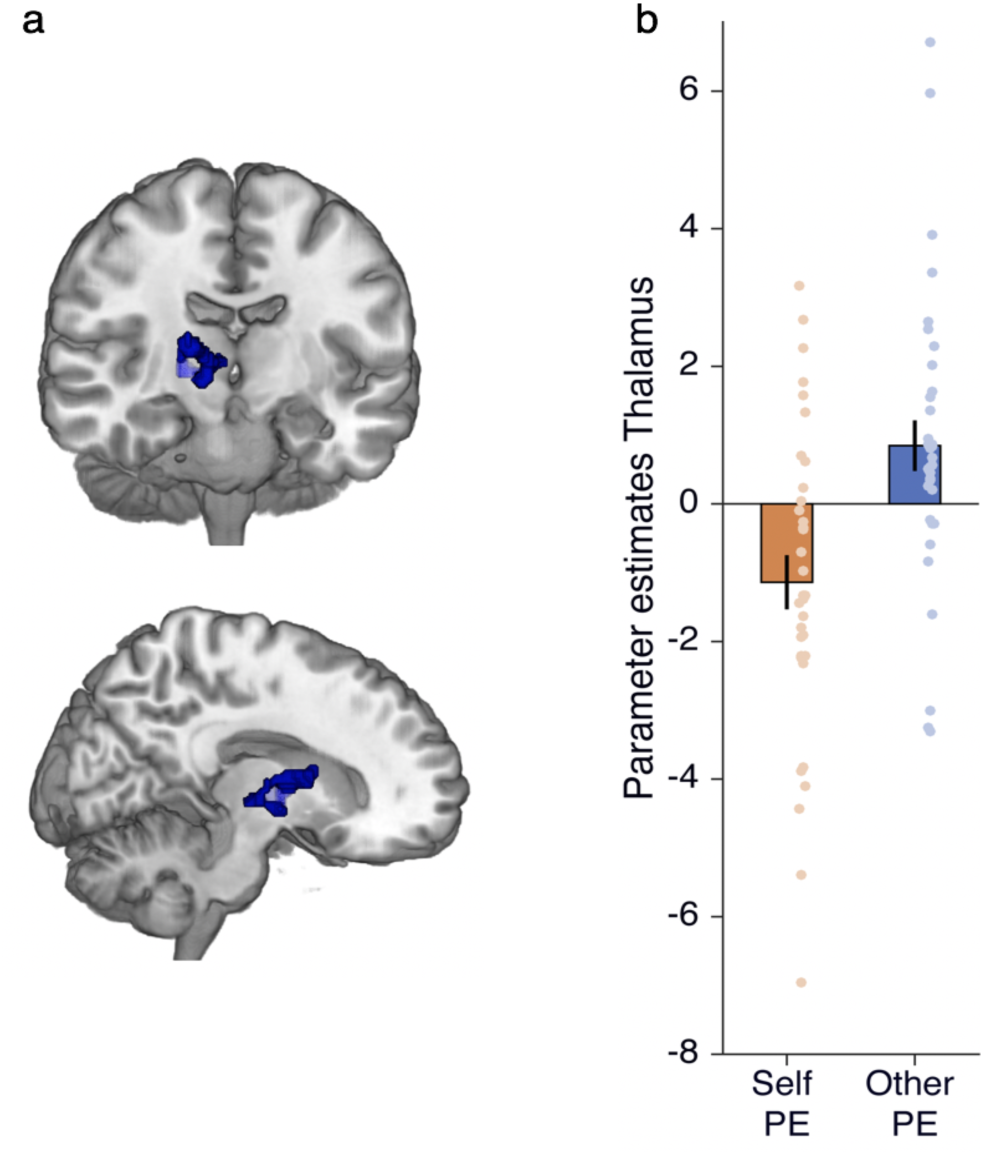
Thalamus/caudate signal distinguishes model-free prediction errors for avoiding harm to other vs. self. (a) Thalamus cluster from the contrast other prediction error > self prediction error (x =16, y = −18, z =0, Z=3.50, k=84, *p*=.033, FWE-SVC after initial thresholding at *p*<.001) overlaid on an anatomical scan to show the extent of activation. (b) for illustration, parameter estimates extracted from the thalamus cluster are shown separately for self and other PE.

### Signatures of model-free influence are encoded in sgACC and TPJ

One signature of model-free learning is a tendency to repeat previously rewarded actions and avoid previously punished actions, regardless of experienced transitions^20^. Such a model-free influence is thought to emerge at the time of choice by activating the reinforcement histories of potential actions and driving selection of the most valuable action in terms of its recent history^70^. In the context of our task, through model-free influence, an action that was unpunished on the previous trial should be prioritized for selection (‘stay’), while an action that was punished on the previous trial should be avoided (‘switch’). Importantly, because model-free learning is insensitive to task structure, this process should occur regardless of whether the transition from the first to the second stage experienced on the previous trial was common or rare.

Therefore, to probe the neural signatures consistent with a model-free influence at the time of choice, and any potential differences between self and other conditions, we examined neural responses during ‘switch’ and ‘stay’ choices at the first stage as a function of the outcome on the previous trial (no pain or pain). We created an additional GLM (GLM2) that modelled the onset of self trials after pain, self trials after no pain, other trials after pain, and other trials after no pain, with stay (−1) and switch (1) coded as parametric modulators of each of these onsets. Thus, our analysis examined differential neural encoding of stay vs. switch decisions on the current trial, as a function of the outcome on the previous trial (pain or no pain) and its recipient (self or other).

Our analysis revealed a signal in sgACC consistent with a model-free influence on ‘other’ trials (x=-2, y=36, z=6, K=498; Z = 3.88, *p*=.028, FWE whole-brain corrected). This region was more active during stay relative to switch choices on the current trial, following a ‘no pain’ outcome on the previous trial, selectively in the ‘other’ condition (see **Figure 5a-b** and Supplementary Analysis for more details). Corroborating the view that this signal reflects a model-free influence, responses in sgACC were positively associated with the model-free x recipient interaction in behaviour (*r*(33) = .36, *p*=.04, **Figure 6a**,**b**) such that participants with the largest sgACC difference between stay and switch following no pain for other also showed the strongest prioritization of model-free learning for others relative to self.

**Figure 5.**
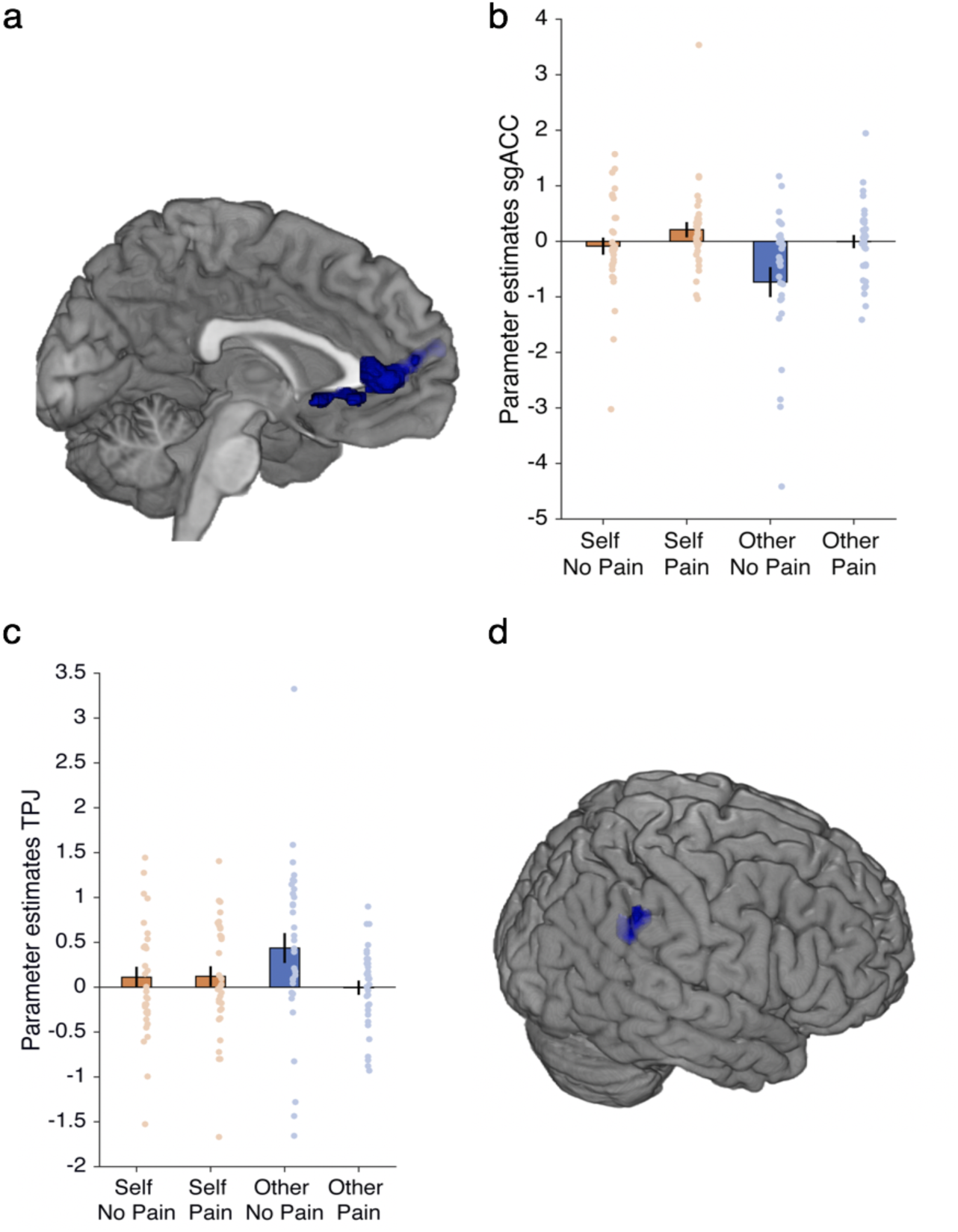
Subgenual anterior cingulate cortex (sgACC) and temporo-parietal junction (TPJ) differentially encode signatures of model-free influence. (a,b) sgACC response (x=-2, y=36, z=6, K=498, *p*=.028, FWE whole brain corrected, after initial thresholding at *p*<.001) to stay vs. switch after no pain for other overlaid on the medial surface of an anatomical scan. The observed BOLD pattern is consistent with model-free behaviour. (c,d) Temporoparietal junction (TPJ) (x=54, y=-38, z=34, Z=3.56, *p*=.03, FWE-SVC after initial thresholding at *p*<.001) tracks switch more than stay after no pain for other. Cluster overlaid on an anatomical scan.

**Figure 6.**
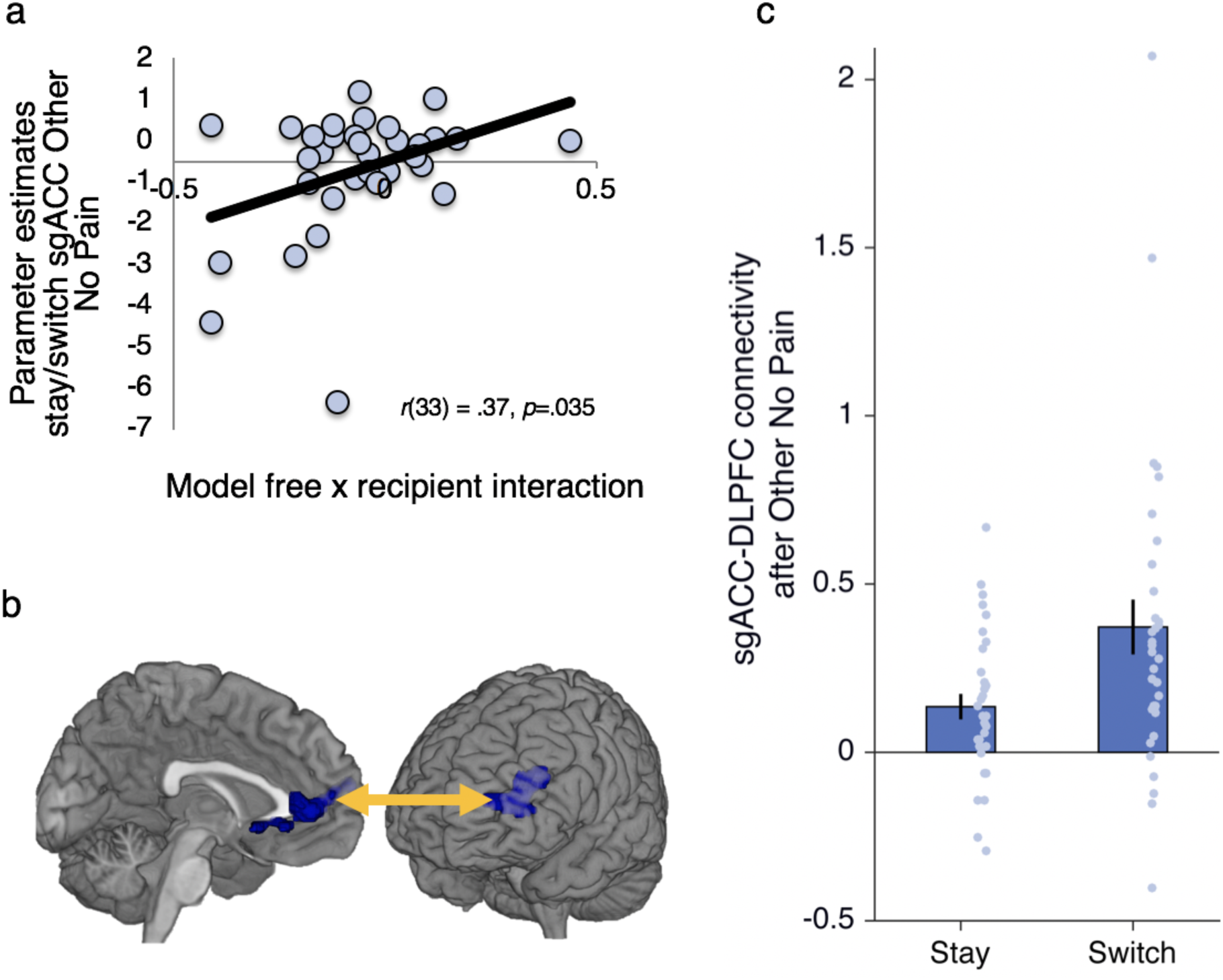
Subgenual anterior cingulate cortex (sgACC) tracks stay vs. switch after no pain for other and connects more strongly to dlPFC when switching after no pain for other. (a) Bivariate association between parameter estimates for stay vs. switch after no pain for other in sgACC and greater model-free behaviour for other (more negative on x means relatively more model-free for other compared to self). (b) sgACC response (x=-2, y=36, z=6, K=498, *p*=.028, FWE whole-brain corrected after initial thresholding at *p*<.001) to stay vs. switch after no pain for other overlaid on the medial surface of an anatomical scan. The observed BOLD pattern is consistent with model-free behaviour. sgACC cluster connects to dorsolateral prefrontal cortex (dlPFC) (x=-46, y=38, z=26, k=382, Z=4.12, *p*=.039, FWE whole-brain corrected after initial thresholding at *p*<.001) during decisions to switch relative to stay after no pain for other. (c) Average slope estimate across participants shows stronger connectivity during switch decisions than stay decisions after receiving no pain for other between sgACC and dlPFC. Switching after no pain goes against a model-free influence.

We observed the inverse pattern (i.e., opposite to what would be predicted by model-free influence) in right TPJ (x=54, y=-38, z=34, Z=3.56, K=39, *p*=.03, FWE-SVC). This region was more active during switch relative to stay choices on the current trial, following a ‘no pain’ outcome on the previous trial, in the ‘other’ condition specifically (**Figure 5c-d** and Supplementary Analysis).

### Increased functional connectivity between sgACC and dorsolateral prefrontal cortex when resisting model-free influence

Behavioral analyses indicated that participants on average showed a mixture of model-based and model-free strategies, but prioritized model-free learning when avoiding harm to others. Thus, we next sought to identify regions that might modulate the model-free effects observed specifically in the ‘other’ condition in sgACC (**Figure 6a**). We therefore conducted psycho-physiological interaction (PPI) analyses (GLM3) to assess functional connectivity between sgACC and the whole brain as a consequence of staying versus switching after no pain for other (see methods for additional details of analyses). This analysis identified a significant negative association between activity in sgACC and dorsolateral prefrontal cortex (x=-46, y=38, z=26, k=382, Z=4.12, *p*=.039, FWE-whole brain corrected after initial thresholding at *p*<.001).

To understand the nature of this effect we plotted the average slope of sgACC and dlPFC connectivity during stay and switch conditions (**Figure 6b**,**c**). This showed that there was a stronger positive coupling between sgACC and dlPFC during switch choices compared to stay choices after receiving no pain for another person (**Figure 6b-c**). We did not observe significant coupling between sgACC and dlPFC during switch or stay choices following no pain for self (Supplemental Analyses).

### Individual differences in moral judgment relate to model-free moral learning

Finally, we investigated whether individual differences in moral judgment were related to individual differences in model-free moral learning and its neural basis. We examined two aspects of moral judgment. First, motivated by theories suggesting that ‘anti-utilitarian’ judgments in moral dilemmas might be driven by model-free processes^3,27^, we measured individual differences in utilitarian reasoning using the Oxford Utilitarianism Scale^71^. Because our study concerned harmful outcomes, we predicted that we would observe correlations with the ‘instrumental harm’ component of the scale, which measures a permissive attitude towards harming one person in order to help many others, but not the ‘impartial beneficence’ component of the scale, which measures an impartial attitude towards helping others. First, we tested whether utilitarianism was correlated with model-free behavior for others (relative to self). We found a significant relationship between instrumental harm and model-free moral behaviour, such that those who were the most anti-utilitarian were the most model-free (*r*(36) = 0.37, *p* = 0.026). However, there was no correlation with the impartial beneficence subscale (*r*(36) = −.059, *p* = 0.731). Next, given past work linking utilitarian reasoning with dLPFC function^72,73^, we predicted that utilitarianism would positively predict dLPFC-sgACC connectivity when resisting model-free influence. This prediction was supported, with dLPFC-sgACC connectivity positively correlated with instrumental harm (*r*(33) = 0.43, *p* = 0.012), but not impartial beneficence (*r*(33) = −.11, *p* = 0.537).

Second, we probed the sensitivity of moral wrongness judgments to how much suffering an action inflicts on a victim (‘outcome sensitivity’), versus how aversive it feels to perform the action (‘action sensitivity’^74^). Past work has connected the former with model-based learning and the latter with model-free learning^3,27^. However, we note that model-free learning is directly sensitive to recent outcomes^18^, which might lead to an association between model-free behavior and outcome sensitivity. Participants evaluated the moral wrongness of 23 harmful actions that varied independently in how much suffering they would cause, versus how aversive they would feel to perform. Action sensitivity and outcome sensitivity were inversely correlated (*r*(36) = −.40, *p* = .016). Partial correlations controlling for action sensitivity revealed that outcome sensitivity was positively correlated with several aspects of model-free moral learning (**Figure 7**), including the tendency to switch following harm to others (*r*(36) = −.37, *p*=.027), the strength of model-free prediction error signals for other vs. self in thalamus/caudate (*r*(34) = .385, *p* = .025), and the strength of model-free influence in sgACC (*r*(33) = −.374, *p* = .032). The reverse partial correlations testing for the effect of action sensitivity whilst controlling for outcome sensitivity were not significant (all *p*’s>.08, all |r’s| <-.30). Together these findings suggest that a natural tendency to engage in model-free moral learning when avoiding harm to others is related to how moral judgments of others’ harmful actions track with harm severity.

**Figure 7.**
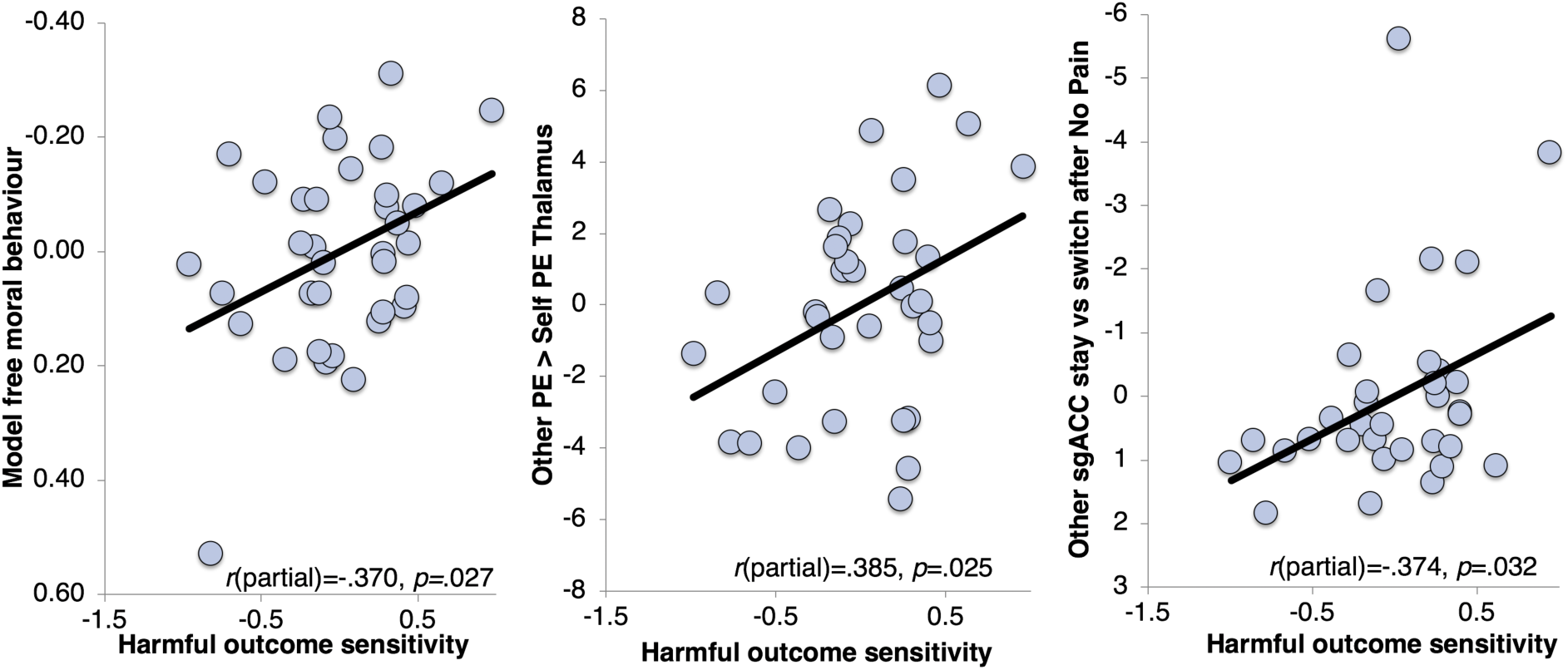
Sensitivity to harm in moral judgments correlates with model-free moral behaviour and its neural correlates. (a) Partial correlation controlling for harmful action sensitivity between harmful outcome sensitivity and model-free moral behaviour. Note the values are reversed on the y axis to depict that greater model-free behaviour is associated with greater harmful outcome sensitivity (b) Partial correlation controlling for harmful action sensitivity, between harmful outcome sensitivity and prediction errors of pain avoidance in the thalamus/caudate for Other compared to Self (c) Partial correlation controlling for harmful action sensitivity between harmful outcome sensitivity and parameter estimates extracted from the parametric regressor for stay vs switch after no pain for other in subgenual anterior cingulate cortex (sgACC). Note that values are reversed on the y axis such that a greater tendency to stay vs switch tracked in sgACC correlates with harmful outcome sensitivity.

## Discussion

Learning to avoid actions that harm other people is a fundamental prerequisite for moral behaviour. Here we show that people prioritize model-free learning when actions have the potential to harm others, and that learning to avoid harming others (vs. self) has a distinct neural signature. The thalamus/caudate differentially encoded prediction errors of pain avoidance for self versus other, whilst sgACC and right TPJ tracked positively and negatively with model-free influence on pain avoidance at the time of choice. Overriding model-free influence when choices affected others invoked stronger connectivity between sgACC and dlPFC. Finally, multiple aspects of moral judgment were associated with model-free moral learning and its neural correlates.

In the context of our study, model-free moral learning manifested as a reduced likelihood of repeating actions that harmed others on the previous trial, regardless of whether such actions typically led to states with a high likelihood of harmful outcomes. Our behavioural finding that people were more model-free when learning to avoid harming others relative to themselves suggests that potentially harmful actions might be prioritized for automatic avoidance as a default. Given the importance of avoiding harm to others for social life, such a learning mechanism would be socially adaptive. Our findings are consistent with prior work showing that harmful social choices are slower than helpful choices^5,51,52^ as well as evidence that morality constrains mental representations of what actions are considered possible, with harmful actions removed from choice sets as a default stance^50^. Repeatedly assigning negative action values to harmful actions over the course of one’s life might automatically remove such actions from consideration, even though in some cases locally harmful actions can lead to wider benefits (e.g., a surgeon cutting open a patient to remove a cancerous tumor). If such a learning strategy is socially adaptive, this raises interesting questions about whether model-free learning can be considered an “optimal” strategy in the ecological sense.

An alternative explanation for our behavioral findings is that model-based learning is effortful^26^, and people choose to put in less effort to benefit others^75^. However, this explanation seems unlikely given that model-free moral learning in our study was specifically driven by a lower probability of repeating choices that harmed others. In other words, participants were more likely to actively switch their choices following harm to others, rather than passively sticking with the status quo. Our results were also not due to differences in the subjective perception of the harmfulness or aversiveness of the outcomes for others compared to self, as participants overall rated shocks received for others in the task as being just as aversive as shocks received for themselves. Furthermore, we did not observe differences in choice consistency (captured by the temperature parameter in our model) during learning for self vs. others. Since choice consistency is related to task engagement, if our behavioral effects reflected reduced effort for others, we would expect to see lower choice consistency when learning for others than self. Together these alternative considerations suggest that people might naturally prioritize model-free strategies/behaviours when avoiding harm to others, and these effects may not simply be explained by less effort or engagement when actions affect others relative to oneself.

Turning to the neural findings, we observed a signal in the thalamus, extending to the caudate, that differentially encoded model-free prediction errors when learning to avoid harming others versus self. These subcortical regions were previously observed to encode value during moral decisions to avoid profiting from others’ pain^4^ and play a critical role in associative learning and moral-decision-making more broadly^40,42,44,45^. The thalamus is often linked to the processing of the affective dimension of pain in addition to its sensory properties^76^. For example, microstimulation of the thalamus can invoke affective memories of previously experienced pain^45^. The thalamus/caudate signal differed from the adjacent ventral striatum response that positively tracked model-free prediction errors regardless of the recipient of the outcome, consistent with a previous study using a similar task with rewarding outcomes for self only^20^. These findings suggest that multiple sub-cortical areas support model-free moral learning, perhaps with ventral striatum providing a generic model-free prediction error signal that is insensitive to outcome valence and outcome recipient, and thalamus/caudate providing additional information about social context.

Another signature of model-free influence was observed in sgACC at the time of choice, contingent on the outcome of the previous trial and specific to the ‘other’ condition. Specifically, signal in sgACC was higher when participants repeated actions that previously avoided harming others, but not during similar choices for oneself. This pattern is consistent with model-free behavior, and individual differences in model-free behavior tracked with individual differences in sgACC response at the time of choice. Notably, previous work has implicated sgACC in model-free learning to gain rewards for others but not self^53^ and in receiving unexpected positive feedback from others^77^, suggesting this region might compute learning signals that are specific to social settings. More broadly, activity in sgACC has been positively associated with prosocial and moral behaviours^15,54,55,57^.

Collectively these findings suggest that sgACC might bias decision-making away from choices that could harm others. However, this default strategy is not always appropriate, for instance in settings where a typically harmful action might lead to a better outcome. We observed signals at the time of choice in two areas that negatively tracked with model-free influence. The first, in right TPJ, showed a response pattern precisely opposite to that observed in sgACC. Signal in rTPJ was higher when participants abandoned a choice that previously avoided pain for others, but not during similar choices for oneself. In addition, overriding model-free influence at the time of choice was associated with increased functional connectivity between sgACC and dLPFC. These two regions showed stronger coupling on trials where participants abandoned a choice that previously avoided pain for others, compared with trials where participants repeated actions that previously spared others from pain. Although we cannot confidently attribute these patterns to model-based control, past work has implicated dLPFC and TPJ in model-based learning and decision-making^31,70^. One intriguing possibility is that sgACC promotes social harm avoidance as a default, while TPJ and dLPFC provide contextual information that enables this default to be overridden when appropriate. Research implicating these regions in the adjustment of moral decisions to blame and punishment provides initial support for this possibility^5,32,33,39,47^.

Finally, we observed correspondences between model-free learning, its neural substrates, and moral judgments. Theoretical work has proposed links between model-based/model-free learning and moral judgment^3,27,28,78^, but empirical support for such links has been scarce. We probed two aspects of moral judgment. First, we examined individual differences in the dimension of ‘utilitarian’ moral reasoning that justifies harming one person to help many others. Consistent with our predictions, as well as work highlighting a link between dLPFC activity and ‘utilitarian’ judgments^73^, we found a positive relationship between instrumental harm and dLPFC-sgACC connectivity when resisting model-free influence. We also observed that those who were the most anti-utilitarian were the most model-free.

Second, using a task that asks participants to judge how morally wrong it would be to perform a series of violent actions that varied independently in terms of how much suffering they would inflict (‘outcome sensitivity’), versus how aversive they felt to perform (‘action sensitivity’)^74^. We found that individual variability in both the behavioural and neural signatures of model-free learning was specifically correlated with outcome sensitivity, but not action sensitivity. Those people whose moral wrongness judgments were more sensitive to the severity of harmful outcomes were less likely to repeat decisions that harmed others, and showed stronger model-free prediction error signals in the thalamus and caudate, and stronger responses in sgACC when repeating decisions that previously avoided harming others. Model-free learning has previously been suggested to explain why actions that typically harm others feel aversive to perform, even when they are not actually harmful^3,27^. Thus, one might expect that a greater tendency to engage in model-free moral learning should predict action sensitivity in moral judgments, rather than outcome sensitivity. However, model-free learning is directly sensitive to recent outcomes^18^, and in the context of our task, manifested as a tendency for choices to be immediately sensitive to harmful outcomes for others. Thus, individual differences in sensitivity to others’ harm could be commonly associated with model-free moral learning and outcome sensitivity in moral judgments. Overall, these findings provide preliminary evidence linking model-free learning to individual differences in moral judgments, effects that could be investigated more extensively in larger samples.

More broadly, our findings highlight differences in the neurocognitive mechanisms engaged in learning to avoid harming oneself versus others that underscore the unique demands of social decision-making. One important feature of decisions that affect others (as opposed to oneself) is that it is far more difficult to build a model that incorporates others’ preferences and beliefs than a model that captures only one’s own preferences. Furthermore, it is impossible to evaluate the accuracy of such models, given that the subjective experiences of others are fundamentally unknowable^79,80^. Utilitarian approaches to moral decision-making that involve maximizing well-being for all sentient beings^81^ may thus be computationally intractable for the model-based system. Rule-based approaches, like those enshrined in deontological theories of morality^82^ circumvent the need for complex model building and may be socially adaptive even for simple social decisions like the ones studied here. Whether the prioritization of model-free learning extends to other kinds of social decisions, such as acting to obtain rewards for others, avoiding monetary losses or indeed even social decision-making in nonhuman species, is an important topic for future study.

Overall, we observed that when learning to avoid harm to others (versus self), participants showed a stronger relative balance toward model-free over model-based learning. Multiple model-free learning signatures were apparent in behaviour as well as cortical and subcortical areas that distinctly process harm avoidance for others compared to self. These findings could have important implications for theories of learning and moral-decision-making as well as disorders associated with impaired avoidance learning and social cognition.

## Methods

### Experimental procedures

#### Participants

Forty-one right-handed healthy adults were recruited through university participant databases. Exclusion criteria included previous or current neurological or psychiatric disorder, non-normal or non-corrected to normal vision, previous participation in studies involving social interactions and/or electric shocks and contraindications that prohibited MRI scanning. One participant was excluded as they reported that they did not believe their decisions would affect another person in the post-scanning debrief. Three participants were excluded because the logistic regression analysis of their choice behaviour did not converge. In two of these participants, this was because they had less than 5% switch trials; in the third participant, there was a very high correlation (>0.8) between the outcome and transition regressors on switch trials. One participant was excluded from the fMRI analysis due to distortions in the scan caused by metal artefacts (braces) and another for excessive head motion. This left a final sample of 36 participants for behavioural analyses (16 female, age 18-36) and 34 participants (16 female, age 18-36) for the parametric fMRI analyses. For the stay/switch analysis one further participant was excluded for having no variance in at least one regressor making their fMRI GLM inestimable. With 33 subjects we had 80% power to detect a ‘medium’ effect size of *d* = 0.50 at alpha = 0.05 (two-tailed), an effect size smaller than typically reported in this field, indicating sufficient power. All participants gave written informed consent and the study was approved by the University of Oxford Medical Sciences Division Ethics Committee.

#### Procedure

Participants completed a pain thresholding procedure which was based on previous studies of self and other pain processing^4,5^. The pain thresholding procedure allowed us to control for heterogeneity of skin resistance between participants to ensure the delivered shocks would be rated at a matched subjective level of pain intensity and also to provide participants with full experience of the shocks before the learning task to ensure their choices were truly guided by knowledge of the pain and no pain outcomes. Participants were then assigned to roles of either ‘decider’ or ‘receiver’ using a role assignment procedure that has been used in several previous studies (see^5,75^). Briefly, participants were instructed to wear coloured rubber gloves to hide their identity. They then stood either side of a door and waved to one another so that they knew another person was there but could not discern any information about the other participant’s age or gender. Next a coin was flipped to decide who would draw a ball out of a box first and then each participant drew a ball. The experimental participant was then told that the colour of their ball meant they had been assigned to the role of decider. Before completing the task in the scanner participants performed a practice task of one block that did not specify whether outcomes were for self or other and they were told no actual shocks would be received during the practice. This practice task, which used stimuli not used in the experimental task, allowed participants to become familiar with the transition structure of the task, that they were told would remain the same in the main experiment. The main experiment consisted of 4 blocks of 68 trials (136 trials for self and 136 trials for other) and lasted ∼45 minutes.

#### Experimental task

We adapted features from two variants of a task designed to distinguish model-free and model-based learning^20,26^ (**Figure 1, Supplemental Experimental note**). Participants were presented with two fractal images that probabilistically led them to one of two ‘states’ where they were required to make a second choice that was followed by a symbol indicating the receipt of pain or the receipt of no pain (neutral). At the beginning of each block and on each trial, they were told whether they were playing for themselves (‘You’) meaning the painful outcomes would be delivered to themselves, or for the other participant (‘Receiver’), indicating that the painful outcomes would be delivered to the other participant. In order to match the self and other conditions in terms of pain stimulation, no electric shocks were delivered during the scan. However, participants were told that 10% of the electric shocks that they acquired during the task would be given to themselves at the end of the session and 10% of the electric shocks they accumulated for the other participant would be delivered to the other participant at the end of the scanning session. In order to account for potential differences in pain perception, participants were instructed that for ‘You’ trials we would use the voltage setting that corresponded to *their* level 8 rating, and for the “Receiver” trials, we would use the voltage setting that corresponded to *the Receiver’s* level 8 rating (full instructions can be downloaded at (OSF https://osf.io/3stp9/files/). At the end of the scan participants also rated how they felt when obtaining a shock for themselves and the receiver during that task on a 0-10 point scale from ‘very negative’ (0) to ‘very positive’ (10). Participants perceived the shocks to be aversive for both self (*M*=3.38, *SD* = 1.44) and other (*M*=3.30, *SD* = 1.61) (*t*(36) = .279, *p*=.782) suggesting that the perception of harm was equivalent.

Participants were instructed that the probability of reaching each of the two ‘states’ would remain fixed throughout the task but the probability that each stimulus would deliver pain or no pain would change throughout the task so that they needed to keep on learning. Adding the ‘receiver’ trials meant that fewer trials, overall, were included in each condition compared to the original paradigm^20^. We therefore made two modifications to the original paradigm in order to better sample variability in behaviour. We bounded the probability that the four second-stage options would deliver pain between 0 and 1 (instead of 0.25 and 0.75 in^20^) and we increased the drift rate of the probabilities to 0.2 (instead of 0.025 in^20^). Moreover, we specifically chose to implement the main aspects off the Daw et al.^20^, version of the paradigm rather than using all the modifications described in^26^, as the Daw et al.^20^, paradigm allowed us to assess peoples’ relative balance between model-free or model-based behaviour^20,26^ when there was no particular incentive to being more model-free or model-based on the task because the degree of model-basedness did not affect the number of shocks received.

#### Statistical analysis of behavioral data

Analyses of behavioural data were performed in SPSS 25 (Armonk, New York: IBM Corp, for bivariate correlations), R (version 1.1.423, for linear mixed-effects modelling, lme4, version 1.1-21) and Matlab 2015b (for logistic regression analyses on the probability of stay). For Fig 2a, we calculated the % of stay choices after common or rare transitions following pain or no pain outcomes (2×2). For regression analyses using lme4 in R, we coded Stay as 1 and Switch as 0 and created regressors for transition type, outcome, outcome x transition type, agent x outcome and agent x outcome x transition type. We used Bound Optimization by Quadratic Approximation (bobyqa) with 1e5 function evaluations. We examined bivariate associations between the interactions with agent from the regression analyses and neural responses with individual differences in utilitarianism using the Oxford Utilitarianism Scale (OUS)^71^ and action and outcome sensitivity from the Harmful Action Outcome Scale^74^.

#### Moral judgment measures

Participants completed the OUS instrumental harm (OUS-IH), OUS Impartial Beneficence (OUS-IB) scales and the HAO scales via an online link in the preceding weeks before the scanning session. The OUS-IH consists of 4 items reflecting a relative willingness to cause harm to others in order to bring about the greater good (e.g., “It is morally right to harm an innocent person if harming them is a necessary means to helping several other innocent people”). The OUS-IB subscale consists of 5 items reflecting endorsement of the impartial maximization of the greater good even at a cost to oneself (e.g. if the only way to save another person’s life during an emergency is to sacrifice one’s own leg, then one is morally required to make this sacrifice). Participants rated these items on a 7-point scale (*1 = strongly disagree, 7 = strongly agree).* A mean score was then computed for all participants.

For the HAO participants were presented with a scenario about two people, Carl and John. They are told John is terminally ill and sincerely wants to die and has asked another person, Carl, to perform a mercy killing. Participants then rated how morally wrong each of the 23 methods of killing were on a scale from 1 – the least morally wrong, to 10 – the most morally wrong. In a previous study^74^ participants were asked to rate the action value and outcome value of these different methods of killing. To assess action value, the researchers asked participants to rate how *upsetting* it would be to “act out” performing each behavior as though it were part of a movie script. To assess outcome value, they asked them to rate how much *suffering* each act would impose. We used the mean action and outcome scores derived from this initial paper to predict the ‘wrongness’ scores in our current sample. This analysis created two different beta weights for each participant corresponding to the action and outcome sensitivity, respectively.

#### Computational modelling of behavioural data

For modelling of choice behaviour using trial-by-trial updates, we proceeded in two steps. First, we evaluated a number of models separately based on their performance on the self and other blocks. This was to probe whether the same model would win for self and other blocks, allowing us to rule out participants might employ entirely different strategies in the two block types. The following models were fitted:

<u>(1) 7-parameter: full model specified by Daw et al. using parameters:</u> learning rates for stage 1 and 2 (αS1, αS2), temperature parameters for stage 1 and 2 (βS1,βS2), a perseverance parameter (ρ), an eligibility trace (λ) and a model-free/based weighting parameter (ω); for full details of model see Supplementary information and^20^

(2) 6-parameter model, as (1) but with λ=1 (λ was shown to have a high mean value and small variance in previous work e.g.^62^)

(3) 5-parameter model, as (1) but with only one α and β for stage 1 and 2

(4) 4-parameter model, as (3) but with λ=1

(5) 5-parameter model, as (4) but with two learning rates for pain and no pain outcomes (αPain, αNoPain)

Models were fitted using a hierarchical Bayesian model fitting approach described in detail in^83,84^. It finds the maximum *a posteriori* estimate of each parameter for each subject using a prior distribution for each parameter which helps to regularise and constrain parameters. The algorithm uses Expectation-Maximization (EM)^85^ and parameters were transformed to a logistic or exponential distribution to enforce constraints and ensure normality such that 0<{α,ω}<1, {β,λ}>0.

For formal model comparison, we report the Bayesian Information Criterion (BIC) based on the log-likelihood, and computed the model evidence by integrating out the free parameters (BICint^83,84^; **Supplementary Tables 2 and 4**). Exceedance probabilities were calculated by feeding the BICint into SPM’s function spm_BMS (http://www.fil.ion.ucl.ac.uk/spm/software/spm8).

The five-parameter model with separate learning rates for pain and no-pain outcomes best explained behaviour in both self and other conditions (**Supplementary Table 2**). We report the difference between the best-fitting parameters, but this method has a caveat. Because of the nature of hierarchical fitting, which uses separate priors for self and other parameters, this method is somewhat biased towards finding differences. Meanwhile, fitting self and other parameters using the same priors, is overly conservative and biased against finding differences.

To resolve whether the parameter ω differed between self and other blocks, in line with results from the basic logistic regression analyses, and without introducing any such biases, in a second step, we therefore fitted three models to the merged data of both self and other blocks:

1. 5-parameter model (as (5) above) with all parameters shared between self/other
2. 6-parameter model with αPain, αNoPain, β and ρ shared but ω split into ωSelf and ωOther
3. 7-parameter model with αPain, αNoPain, β shared and ρ and ω split into ρSelf, ρOther, ωSelf and ωOther

As described above, model comparison was performed based on BICint values (**Supplementary Table 4**). The mean parameter estimates are shown in **Supplementary Fig 3** and **Supplementary Table 5**. We also simulated data from our participant schedules and showed that we had reliable parameter recovery^86^

#### FMRI acquisition and analysis

Multiband T2*-weighted echo planar imaging (EPI) volumes with blood oxygenation-level–dependent (BOLD) contrast were acquired using a Siemens Prisma 3T MRI scanner. The EPI volumes were acquired in an ascending manner, at an oblique angle (≈30°) to the AC-PC line to decrease the impact of susceptibility artefacts in the orbitofrontal cortex. We used the following parameters. Voxel size 2 × 2 × 2, TE=30 ms; repetition time=1570ms; flip angle=90°; field of view=216 mm. The structural scan was acquired using a magnetization prepared rapid gradient echo (MPRAGE) sequence with 192 slices; slice thickness=1 mm; TR=1900 ms; TE=3.97 ms; field of view=192 mm x 192mm; voxel size=1×1×1 mm resolution.

Fmri data were analysed using SPM12 (www.fil.ion.ucl.ac.uk/spm). Images were realigned and unwarped using a fieldmap and co-registered to the participant’s own anatomical image. The anatomical image was processed using a unified segmentation procedure combining segmentation, bias correction, and spatial normalization to the MNI template using the New Segment procedure; the same normalization parameters were then used to normalize the EPI images. Lastly, a Gaussian kernel of 8 mm FWHM (SPM default) was applied to spatially smooth the images.

Before the study, example first-level design matrices were checked to ensure that estimable GLMs could be performed with independence between the parametric regressors: value difference at the first-stage choice, the state prediction error at stage 2, and a model-free prediction error at the time of the outcome. This allowed us to look at value and prediction error responses independent from one another. We also tested a GLM that coded switch vs. stay trials as a parametric modulator at the time of choice dependent on the previous outcome. Again this GLM could be estimated with independence (See Supplementary Figure 1). We convolved these different event types with SPM’s canonical haemodynamic response function. All events were modelled as stick functions with 0 duration.

For GLM1, each of these regressors was associated with parametric modulators taken from the computational model. At the time of the first stage choice this was the value difference from the hybrid model combining model-free and model based learning. At the time of the second stage choice this was the state prediction error for the transition from stage 1 to stage 2, and at the time of the outcome this was the model free prediction error (since the behavioural differences were in the model-free parameters). In all cases, values were modelled separately for the onsets of self and other trials. As in^20^, we fixed the parameters to the average values for self and other (**Supplementary Table 4**) but allowed ω to vary.

For GLM2 we modelled whether participants stayed or switched at the first-stage choice relative to the outcome on the previous trial, i.e. no pain or pain. Due to the smaller number of trials included in this analysis we coded stay and switch as a parametric regressor with values of 1 assigned to switch and −1 assigned to stay. One participant did not have a trial in at least one of these regressors and was therefore excluded from the stay-switch analysis. For all GLMs in some participants, an extra regressor modelled all missed trials, on which participants did not select one of the first-stage choices.

GLM3 and GLM4 corresponded to our psychophysiological interaction analyses. We defined a seed region in the sgACC using a 6mm sphere based on the peak co-ordinates from our analyses (for completeness we also ran PPI analyses using seed regions in thalamus and TPJ, see Supplementary Text). We then extracted the physiological variable and the psychophysiological interaction terms for stay vs. switch after no pain for other (GLM3) and stay vs. switch after no pain for self as a control analysis (GLM4). These PPI terms were entered into the GLMs along with all previous regressors that specified the events of our study as described above. In all GLMs, six head motion parameters modelled the residual effects of head motion as covariates of no interest. Data were high-pass filtered at 128 s to remove low-frequency drifts, and the statistical model included an AR(1) autoregressive function to account for autocorrelations intrinsic to the fMRI time-series.

Contrast images from the first level were input into flexible-factorial designs. Following standard procedures, main effects are reported at *p* < .05, family-wise error (FWE) cluster corrected across the whole brain after initial thresholding at *p*<.001, or *p* < 0.05 FWE small volume corrected (SVC) after initial thresholding at *p*<.001 for regions where we had a strong *a priori* hypothesis^87,88^. These areas were defined anatomically and included subcortical regions of the caudate and thalamus (taken from the WFU PickAtlas Toolbox), bilateral posterior TPJ (from^89^) the sgACC (areas s24 and 25 from^90^ and the dlPFC (areas 46v and 9 taken from^91^). These ROIs were also used to confirm anatomical labelling. We additionally applied a false discovery rate correction (FDR) for the number of ROI corrections. All ROI comparisons remained significant (*p*< 0.05) when controlling for the number of comparisons using FDR.

## Author contributions

P.L.L, M. K-F and M. J. C designed study. P.L.L and A. A. collected data. P.L.L and M. K-F analysed data. P.L.L, M. K-F, A. A., and M. J. C. wrote paper.

## Acknowledgements

This work was supported by a Medical Research Council Fellowship (MR/P014097/1), a Christ Church Junior Research Fellowship and a Christ Church Research Centre Grant to P. L. L., a Wellcome Trust grant (106164/A/14/Z) and an Academy of Medical Sciences (SBF001\1008) grant to MJC. MCKF was supported by a Sir Henry Wellcome Fellowship (103184/Z/13/Z). The Wellcome Centre for Integrative Neuroimaging is supported by core funding from the Wellcome Trust (203139/Z/16/Z). We would like to thank Ms. Eloise Copland and Dr. Hongbo Yu for assistance with data collection, Dr. Marco Wittman, Dr. Peter Smittenaar, Dr. Quentin Huys, Dr. Elsa Fouragnan, and Dr. Mehdi Keramati for assistance with data analysis, and Dr. Matthew Apps, Prof. Nathaniel Daw, Prof Peter Dayan, Dr Wouter Kool and Ms Mary Montgomery for helpful discussions.

## Declaration of interests

The authors have no competing interests

## Materials and correspondence

Please address correspondence to Patricia L. Lockwood, Miriam C Klein-Flugge, and Molly J Crockett.

## Data availability

All data and code used to generate the figures can be downloaded at: [OSF https://osf.io/3stp9/files/]

Unthresholded statistical maps can be downloaded at: [Identifier to be added upon publication]

