## Supplemental material for "Neural signatures of model-free learning when avoiding harm to self and other"

### Supplementary Methods and Results

### Supplementary Text

#### Experimental note in introduction

When designing our study, we considered multiple variants of two-step tasks that have previously been used in research[1–4]. Our aim here was to specifically design our task to assess people's 'relative balance' between engaging in model-free and model-based strategies using a paradigm that yields equal frequencies of pain versus no pain outcomes independent of the level of model-basedness and thus did not 'push' people towards being more model based. We ultimately opted for a hybrid version of the original Daw et al[4] paradigm and the more recently suggested version by Kool et al[2], including some of the benefits of both of these variants of the task.

We chose to adopt one important feature from the original paradigm by Daw et al.[4] and designed the current paradigm such that there was no incentive to be model-based. In other words, model-free and model-based learning strategies yielded overall similar proportions of positive versus negative outcomes. This meant that participants could not avoid more pain by adopting a model-based over a model-free strategy. This was an important feature for two reasons. First, there is extensive evidence that people value others' outcomes differently from their own[5,6] and are willing to exert more effort to benefit themselves than others[7] If we had adopted a more recent variant of this task by Kool et al. where the model-based strategy more effectively obtained desired outcomes (e.g.,[2]) and observed differences in model-based learning for self vs. other, we would have been unable to rule out the possibility that such differences were due to differential utilities in outcomes for self vs. other. Second, this aspect of the original variant of the task is optimised to measure people's 'relative balance' or 'natural tendency' between engaging in model-free vs. model-based learning. In their paper describing the newer version of the two-step task, Kool et al themselves state that "even though there is no significant relationship between reward and model-based control [in the two-step task], this does not undermine the usefulness of the task for measuring the relative balance of model-based and model-free strategies (page 6-7)." The authors therefore highlighted the usefulness of using a variant more closely based on the original Daw paradigm to assess the relative balance between systems which was the main aim of the current study. By contrast, the newer version[2] focusses on the trade-off between cognitive demand and 'accuracy' (here successfully avoiding harm), which was not the focus of the current study.

However, we also adopted some of the improvements in task design suggested by Kool et al[2]., including changes in drift rate and drifts that include more extreme probabilities to facilitate learning. Specifically, we made the drifting of the random walks faster (0.2 as in Kool, rather than 0.025 in Daw) and bounded the reward(pain) rate between 0-1 rather than 0.25-0.75 as suggested by Kool et al. The first modification allowed us to assess learning within a smaller number of trials (138 per agent) than in Daw et al[42] (201 trials) but comparable to Kool et al[2] (125 trials). The second modification allowed us to do this whilst ensuring the task was not too difficult. Also, because there are currently no studies involving neural data for the Kool et al paradigm we wanted a task that could be used, in part, to replicate the neural findings described in Daw et al.[4].

#### Detailed description of full 7-parameter model

We refer to Daw et al.[4] for a more detailed description of the learning model but repeat a short description with the formulas here for completeness.

We denote trials with *t*, states with *i* (A=1st stage, B/C=2nd stage), actions as *j* and rewards as *r*. Model-free updating was described using simple TD learning, with the only free parameters $\alpha$ denoting the stage 1 and 2 learning rates:

$$Q_{TD}(s_{i,t}, a_{i,t}) = Q_{TD}(s_{i,t}, a_{i,t}) + \alpha_i \delta_{i,t}$$

where the prediction error was defined as

$$\delta_{i,t} = r_{i,t} + Q_{TD}(s_{i+1,t}, a_{i+1,t}) - Q_{TD}(s_{i,t}, a_{i,t})$$

The effect of the eligibility parameter $\lambda$ was to update first-stage actions by second-stage prediction errors

$$Q_{TD}(s_{1,t}, a_{1,t}) = Q_{TD}(s_{1,t}, a_{1,t}) + \alpha_1 \lambda \delta_{2,t}$$

Model-based learning was the same as model-free learning at the second stage. At the first stage, model-based values were modelled as

$$Q_{MB}(s_A, a_j) = P(s_B|s_A, a_j) \max_{a \in \{a_A, a_B\}} Q_{TD}(s_B, a) + P(s_C|s_A, a_j) \max_{a \in \{a_A, a_B\}} Q_{TD}(s_C, a)$$

where $P(s_B|s_{A,}a_A)$ and $P(s_C|s_{A,}a_B)$= 0.7 and $P(s_C|s_{A,}a_A) = 1 - P(s_B|s_{A,}a_A)$ and $P(s_B|s_{A,}a_B) = 1 - P(s_C|s_{A,}a_B)$.

Model-based and model-free action values were arbitrated using the free weighting parameter $\omega$:

$$Q_{net}(s_A, a_j) = \omega Q_{MB}(s_A, a_j) + (1 - \omega) Q_{TD}(s_A, a_j)$$

Finally, the probability of choice was computed using a softmax function with inverse temperature parameters $\beta$ for both stage 1 and 2, and perseverance parameter $\rho$.

$$P(a_{i,t} = a|s_{i,t}) = \frac{\exp(\beta_i[Q_{net}(s_{i,t}, a) + \rho \text{ rep}(a)])}{\sum a'(\exp(\beta_i[Q_{net}(s_{i,t}, a') + \rho \text{ rep}(a')]))}$$

Here, rep(a)=1 if the 1st stage choice was the same as one the previous trial, and 0 otherwise.

**Parameter recovery**

Because schedules had been optimized for the seven-parameter model, but our winning model involved five parameters, with two separate learning rates for pain and no pain outcomes, one inverse temperature parameter and a fixed $\lambda$=1, we tested whether we could recover parameters for this model from simulated data for which we knew the ground truth. For the five parameters ($\alpha$Pain, $\alpha$NoPain, $\beta$, $\rho$, $\omega$), we simulated behaviour using the same schedule given to our participants. We used a wide range of parameter values (total of 4 x 4 x 3 x 3 x 4 = 576 combinations), from a grid of values in the ranges: $\alpha$=[0.2 0.4 0.6 0.8]; $\beta$= [2 3.5 5], $\rho$=[-0.5 0 0.5] and $\omega$= [0.2 0.4 0.6 0.8]. We added noise to each of the five parameters for each simulated agent (from a standard normal distribution multiplied by 0.1) to improve our coverage of possible parameter values. After having generated the behaviour, we refitted the simulated behaviour using fminunc in Matlab. We used the best fit from 10 random starting configurations to avoid local minima.

The correlations between the true simulated and fitted parameter values were: $\alpha$Pain, $\alpha$NoPain= 0.88; $\beta$=0.82; $\rho$=0.96; $\omega$= 0.8. Thus, parameter recovery was reliable for all parameters.

**Switch/stay additional analyses**

In the switch-stay analysis we identified sgACC/TPJ activity after receiving no pain for other (Figure 5). There were no significant results at the whole brain level or in any of our ROIs after no pain for self, or pain for either agent. Due to the coding of switch-stay as a parametric modulator we were unable to statistically assess a full 2 (self/other) x 2 (switch/stay) x 2 (no pain/pain) interaction. However, in sgACC, we additionally confirmed using post-hoc t-tests that there was indeed no response for these three parametric modulators (self no pain ($t(32)$=-.56, $p$=.58), self pain ($t(32)$=1.52, $p$=.14), other pain ($t(32)$=-.05, $p$=.96)). This was also true for right TPJ: it responded to stay more than switch after no pain for other but there was no significant effects in any of the other three conditions (self no pain ($t(32)$=.95, $p$=.35), self pain ($t(32)$=1.09, $p$=.29), other pain ($t(32)$=-.065, $p$=.95)).

**Psychophysiological Interaction analyses**

For completeness we also ran an additional set of GLMs that used seed regions in brain areas apart from sgACC that showed specificity of model-free processing for other compared to self (Thalamus and TPJ). We defined a seed region in these two areas using a 6mm sphere based on the peak co-ordinates from our analyses. We then extracted the physiological variable and the psychophysiological interaction terms for other prediction error > self prediction error (peak in thalamus/caudate) and stay vs. switch after no pain for other. These PPI terms were entered into the GLMs along with all previous regressors that specified the events of our study. In all PPI GLMs, six head motion parameters modelled the residual effects of head motion as covariates of no interest. Neither area showed significant connectivity with any part of the brain at the whole brain or small-volume corrected level.

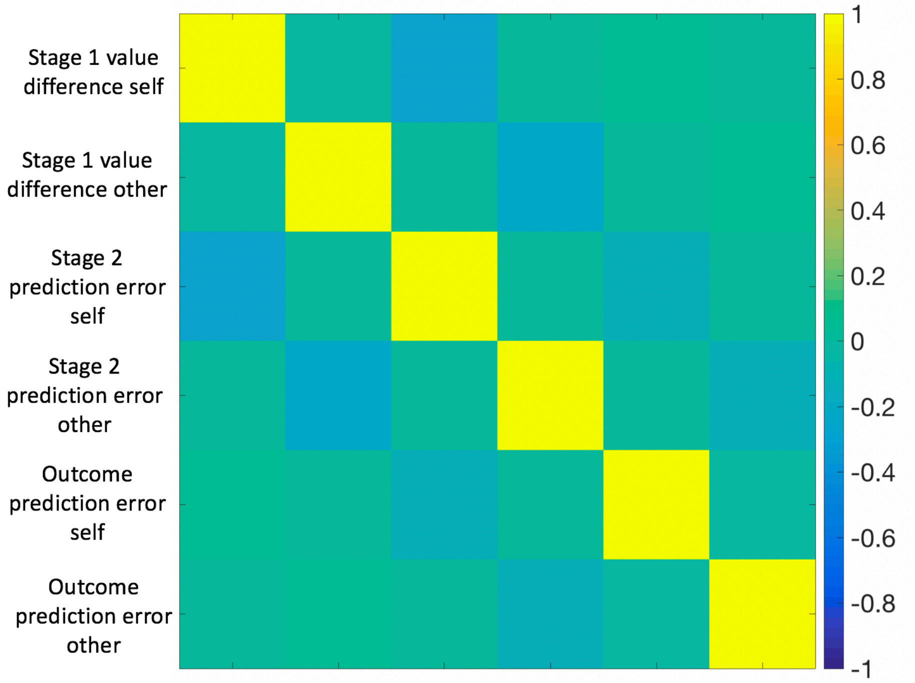


**Supplementary Figure 1**

Correlation between parametric regressors in the model based analyses. All correlations were below r<l0.26l , indicating that conditions could be appropriately estimated with independence from one another.

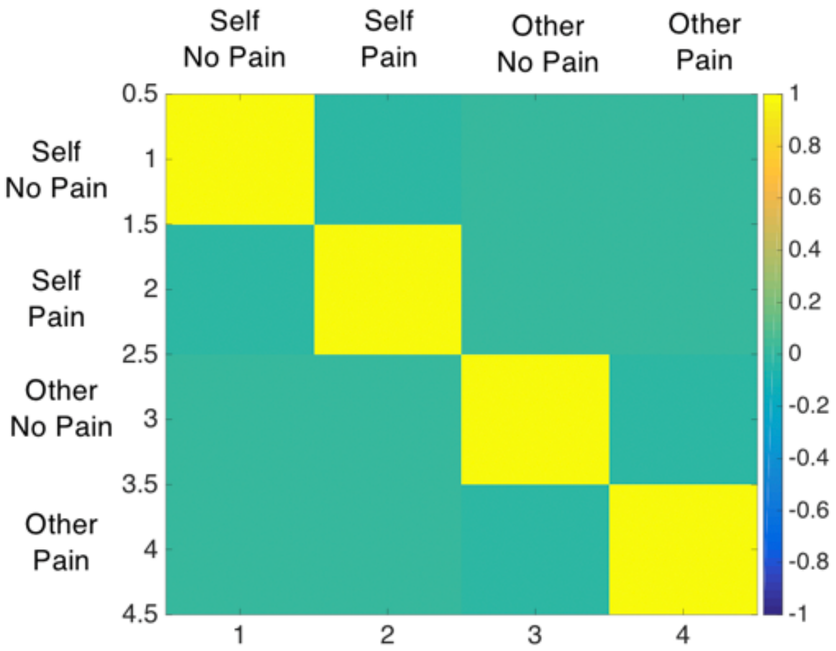


**Supplementary Figure 2.**
Correlation between parametric regressors in switch/stay analysis. All correlations were below *r*<|0.1|, indicating that conditions could be appropriately estimated with independence from one another.

| Brain Region | BA | L/R | Peak voxel | | | k | t | z |
|---|---|---|---|---|---|---|---|---|
| **Main effect: Stage 1 value difference (negative)** | | | | | | | | |
| Anterior cingulate | | L | -6 | 24 | 44 | 625 | 5.69 | 5.11 |
| | | R | 8 | 18 | 46 | | 4.34 | 4.06 |
| Inferior parietral | | L | -34 | -56 | 42 | 1830 | 5.08 | 4.65 |
| | | L | -48 | -40 | 40 | | 4.5 | 4.18 |
| | | L | -26 | -64 | 46 | | 4.37 | 4.08 |
| Inferior parietal | | R | 38 | -54 | 44 | 667 | 4.71 | 4.36 |
| | | R | 42 | -42 | 46 | | 4.28 | 4.00 |
| | | R | 50 | -34 | 50 | | 3.98 | 3.75 |
| Middle Frontal gyrus | | L | -34 | 46 | 22 | 312 | 4.69 | 4.34 |
| | | R | 52 | 22 | 40 | 495 | 4.31 | 4.03 |
| | | R | 48 | 32 | 22 | | 4.01 | 3.78 |
| | | R | 40 | 48 | 22 | | 3.99 | 3.76 |
| **Main effect: Stage 2 prediction error (negative)** | | | | | | | | |
| Anterior cingulate | | L | -6 | 10 | 52 | 906 | 5.34 | 4.85 |
| | | L | -4 | 18 | 44 | | 5.15 | 4.71 |
| | | L | -20 | 4 | 62 | | 3.58 | 3.41 |
| **Main effect: Outcome prediction error (positive)** | | | | | | | | |
| Ventral Striatum | | R | 10 | 12 | -4 | 771 | 8.91 | 7.19 |
| Ventral Striatum | | L | -14 | 8 | -10 | 5994 | 8.5 | 6.96 |
| *ext. Superior frontal gyrus* | | | -18 | 26 | 60 | | 6.3 | 5.55 |
| *ext. Inferior frontal gyrus* | | | -46 | 4 | 26 | | 6.06 | 5.38 |
| Inferior parietal lobe | | L | -48 | -44 | 44 | 4740 | 7.99 | 6.66 |
| | | | -46 | -50 | 54 | | 6.87 | 5.95 |
| *ext. Superior parietal lobe* | | | -36 | -68 | 52 | | 5.92 | 5.28 |
| Superior temporal gyrus | | L | -56 | -38 | -16 | 774 | 6.45 | 5.66 |
| | | | -62 | -24 | 2 | | 3.97 | 3.75 |
| *ext. Inferior temporal gyrus* | | | -46 | -58 | -10 | | 3.41 | 3.26 |
| Precuneus | | R | 30 | -66 | 38 | 2255 | 5.65 | 5.09 |
| | | | 44 | -42 | 52 | | 4.59 | 4.26 |
| | | | 48 | -38 | 46 | | 4.49 | 4.18 |
| Middle occipital gyrus | | R | 32 | -86 | 14 | | 4.32 | 4.04 |
| | | | 30 | -94 | 4 | | 4.31 | 4.03 |

**Supplementary Table 1**. Whole brain analysis ($p$<.05 cluster correction after thresholding at p<.001).

**Table 2 Model comparison self/other separately**

| | Learning rate | Softmax temperature | Perse-verance | Lambda | Model-free model-based weight | negLL | AIC | BIC | BICint | XP | Pro-tected XP |
|---|---|---|---|---|---|---|---|---|---|---|---|
| ***SELF*** | | | | | | | | | | | |
| M5 (5-param) | $\alpha_{Pain}$, $\alpha_{NoPain}$ | $\beta$ | $\rho$ | $\lambda=1$ | $\omega$ | 5342 | 11043 | 11567 | 11143 | 0.9999 | 0.9859 |
| | | | | | | $\Delta$M5 | $\Delta$M5 | $\Delta$M5 | $\Delta$M5 | | |
| M4 (4-param) | $\alpha$ | $\beta$ | $\rho$ | $\lambda=1$ | $\omega$ | 113 | 154 | 49 | 136 | 0.0000 | 0.0035 |
| M3 (5-param) | $\alpha$ | $\beta$ | $\rho$ | $\lambda$ | $\omega$ | 109 | 218 | 218 | 143 | 0.0001 | 0.0036 |
| M2 (6-param) | $\alpha_{Stage1}$, $\alpha_{Stage2}$ | $\beta_{Stage1}$, $\beta_{Stage2}$ | $\rho$ | $\lambda=1$ | $\omega$ | 93 | 257 | 362 | 190 | 0.0000 | 0.0035 |
| M1 (7-param) | $\alpha_{Stage1}$, $\alpha_{Stage2}$ | $\beta_{Stage1}$, $\beta_{Stage2}$ | $\rho$ | $\lambda$ | $\omega$ | 75 | 293 | 503 | 194 | 0.0000 | 0.0035 |
| ***OTHER*** | | | | | | | | | | | |
| M5 (5-param) | $\alpha_{Pain}$, $\alpha_{NoPain}$ | $\beta$ | $\rho$ | $\lambda=1$ | $\omega$ | 5318 | 10995 | 11520 | 11089 | 0.9588 | 0.6752 |
| | | | | | | $\Delta$M5 | $\Delta$M5 | $\Delta$M5 | $\Delta$M5 | | |
| M4 (4-param) | $\alpha$ | $\beta$ | $\rho$ | $\lambda=1$ | $\omega$ | 109 | 146 | 41 | 153 | 0.0001 | 0.0748 |
| M3 (5-param) | $\alpha$ | $\beta$ | $\rho$ | $\lambda$ | $\omega$ | 92 | 184 | 184 | 157 | 0 | 0.0747 |
| M2 (6-param) | $\alpha_{Stage1}$, $\alpha_{Stage2}$ | $\beta_{Stage1}$, $\beta_{Stage2}$ | $\rho$ | $\lambda=1$ | $\omega$ | 56 | 183 | 288 | 165 | 0 | 0.0748 |
| M1 (7-param) | $\alpha_{Stage1}$, $\alpha_{Stage2}$ | $\beta_{Stage1}$, $\beta_{Stage2}$ | $\rho$ | $\lambda$ | $\omega$ | 43 | 229 | 439 | 158 | 0.0411 | 0.1005 |

**Supplementary Table 2**. Separate model comparisons for self and other blocks. For both agents, a model with separate learning rates for no pain and pain ($\alpha$Pain, $\alpha$NoPain), a single temperature parameter ($\beta$), a perseverance parameter ($\rho$) and a model-free/based weighting parameter ($\omega$) performs best (model 5=M5).

| **Table 3** | **Average parameter estimates self/other separately** | | | | | | | | | |
|---|---|---|---|---|---|---|---|---|---|---|
| | **M5** | | **M4** | | **M3** | | **M2** | | **M1** | |
| | **Self** | **Other** | **Self** | **Other** | **Self** | **Other** | **Self** | **Other** | **Self** | **Other** |
| $\alpha/\alpha_{Pain}/\alpha_{Stage1}$ | 0.29 | 0.36 | 0.31 | 0.38 | 0.31 | 0.38 | 0.39 | 0.53 | 0.30 | 0.44 |
| $\alpha_{NoPain}/\alpha_{Stage2}$ | 0.31 | 0.37 | | | | | 0.30 | 0.37 | 0.32 | 0.41 |
| $\beta/\beta_{Stage1}$ | 4.55 | 3.85 | 4.25 | 3.42 | 4.24 | 3.59 | 5.06 | 4.72 | 6.29 | 5.02 |
| $\beta_{Stage2}$ | | | | | | | 3.66 | 4.11 | 3.56 | 4.21 |
| $\rho$ | 0.66 | 0.56 | 0.70 | 0.65 | 0.70 | 0.64 | 0.68 | 0.63 | 0.66 | 0.62 |
| $\lambda$ | | | | | 0.95 | 0.81 | | | 0.53 | 0.54 |
| $\omega$ | 0.59 | 0.46 | 0.61 | 0.57 | 0.59 | 0.44 | 0.63 | 0.65 | 0.48 | 0.53 |

**Supplementary Table 3**. Average parameter estimates extracted for self/other blocks for the five different models. The winning model M5 showed differences in the perseverance parameter ($\rho$) and the model-free/based weighting parameter ($\omega$) between agents.

**Table 4 Model comparison both agents together**

| | Learning rate | Softmax temperature | Perse-verance | Lambda | Model-free model-based weight | negLL | AIC | BIC | BICint | XP | Pro-tected XP |
|---|---|---|---|---|---|---|---|---|---|---|---|
| ***BOTH AGENTS*** | | | | | | | | | | | |
| M5 (5-param) | $\alpha_{Pain}$, $\alpha_{NoPain}$ | $\beta$ | $\rho$ | $\lambda=1$ | $\omega$ | 10800 | **21960** | **22609** | 22175 | 0.8911 | 0.5566 |
| M6 (6-param) | $\alpha_{Pain}$, $\alpha_{NoPain}$ | $\beta$ | $\rho$ | $\lambda=1$ | $\omega_{Self}$, $\omega_{Other}$ | 10771 | 21974 | 22753 | **22151** | 0.1088 | 0.2435 |
| M7 (7-param) | $\alpha_{Pain}$, $\alpha_{NoPain}$ | $\beta$ | $\rho_{Self}$, $\rho_{Other}$ | $\lambda=1$ | $\omega_{Self}$, $\omega_{Other}$ | **10754** | 22012 | 22921 | 22225 | 0 | 0.1999 |

**Supplementary Table 4**. Model comparison when fitting M5 and variants of it to the pooled data from self and other blocks. This does not support a strong conclusion. Nevertheless, the BICint slightly prefers M6 over the others. This model is identical to M5 but models separate model-free/model-based weights for self and other blocks ($\omega_{Self}$ and $\omega_{Other}$). Crucially, there is a significant difference between the parameter estimates obtained for $\omega_{Self}$ and $\omega_{Other}$ providing further support for including both parameters.

**Table 5 Average parameter estimates both agents together**

| | M5 | M6 | M7 |
|---|---|---|---|
| $\alpha_{Pain}$ | **0.35** | **0.35** | **0.35** |
| $\alpha_{NoPain}$ | **0.36** | **0.35** | **0.35** |
| $\beta$ | **3.68** | **3.81** | **3.81** |
| $\rho/\rho_{Self}$ | **0.65** | **0.63** | **0.66** |
| $\rho_{Other}$ | | | **0.57** |
| $\omega_{Both/Self}$ | **0.53** | **0.55** | **0.56** |
| $\omega_{Other}$ | | **0.45** | **0.44** |

**Supplementary Table 5**. Average parameter estimates extracted for both agents fitted simultaneously for the three different models M5-M7. The winning model M6 contained one perseverance parameter ($\rho$) across both agents but separate model-free/based weighting parameters ($\omega$) which differed significantly from each other.
